# Dangling Ends of Third Strand and Duplex Drive Nucleic Acid Triplex Stabilization through Bimodal Association

**DOI:** 10.1101/2025.11.02.686076

**Authors:** Fangyu Zhou, Yiran Liu, Zhiyu Shu, Yongqiang Wang, Gang Chen

**Affiliations:** School of Medicine, The Chinese University of Hong Kong, Shenzhen (CUHK-Shenzhen), No. 2001 Longxiang Blvd., Longgang Dist., Shenzhen, Guangdong, PR China 518172; Guangdong Basic Research Center of Excellence for Aggregate Science, School of Science and Engineering, The Chinese University of Hong Kong, Shenzhen, Shenzhen, Guangdong, PR China 518172; Division of Chemistry and Biological Chemistry, School of Physical and Mathematical Sciences, Nanyang Technological University, 21 Nanyang Link, Singapore 637371; South China Hospital, Medical School, Shenzhen University, Shenzhen, Guangdong, PR China 518116; Shenzhen Key Laboratory of Innovative Drug Synthesis, The Chinese University of Hong Kong, Shenzhen (CUHK-Shenzhen), Shenzhen, Guangdong, PR China 518172; The Chinese University of Hong Kong, Shenzhen (CUHK-Shenzhen) Futian Biomedical Innovation R&D Center, Shenzhen, Guangdong, PR China 518031

**Keywords:** nucleic acid triplex, triplex-forming oligonucleotide (TFO), nucleation, kinetics, dangling ends, base stacking

## Abstract

Nucleic acid triplexes are crucial structural motifs in gene regulation and biotechnology, yet the kinetic principles governing their formation remain poorly understood. While a stability hierarchy of RNA•DNA-DNA > DNA•DNA-DNA > RNA•RNA-RNA, with no DNA•RNA-RNA triplex forming, is known, the kinetic roles of terminal residues remain poorly understood. Here, we employ bio-layer interferometry (BLI) and circular dichroism (CD) spectroscopy to demonstrate that dangling ends from both the third strand (triplex-forming oligonucleotide, TFO) and the duplex dramatically enhance triplex stability. Kinetic analysis reveals this stabilization is primarily driven by a marked increase in the association rate (*k*□□). Crucially, creating a single-base-pair dangling end at either terminus of the duplex enhanced triplex stability more effectively than blunt ends. For example, DNA TFO dTFO5 binding to d(HP5+TA) was enhanced compared to dHP5, and similarly RNA TFO rTFO5 binding to RNA duplex r(HP5+UA) and DNA duplex d(HP5+TA) showed stronger affinity and faster association than to blunt-ended rHP5 and dHP5. Interestingly, removal of a terminal base pair from the blunt-end duplex, generating a TFO dangling end, also enhances binding affinity and association rate. This indicates that both duplex and TFO dangling ends provide critical nucleation platforms, while blunt-ended terminal triples are dynamic and contribute minimally to stability. Thus, our work establishes that optimal triplex formation requires strategic optimization of both TFO and duplex terminal structures through a fundamental kinetic principle (bimodal nucleation).

**Graphical abstract:** 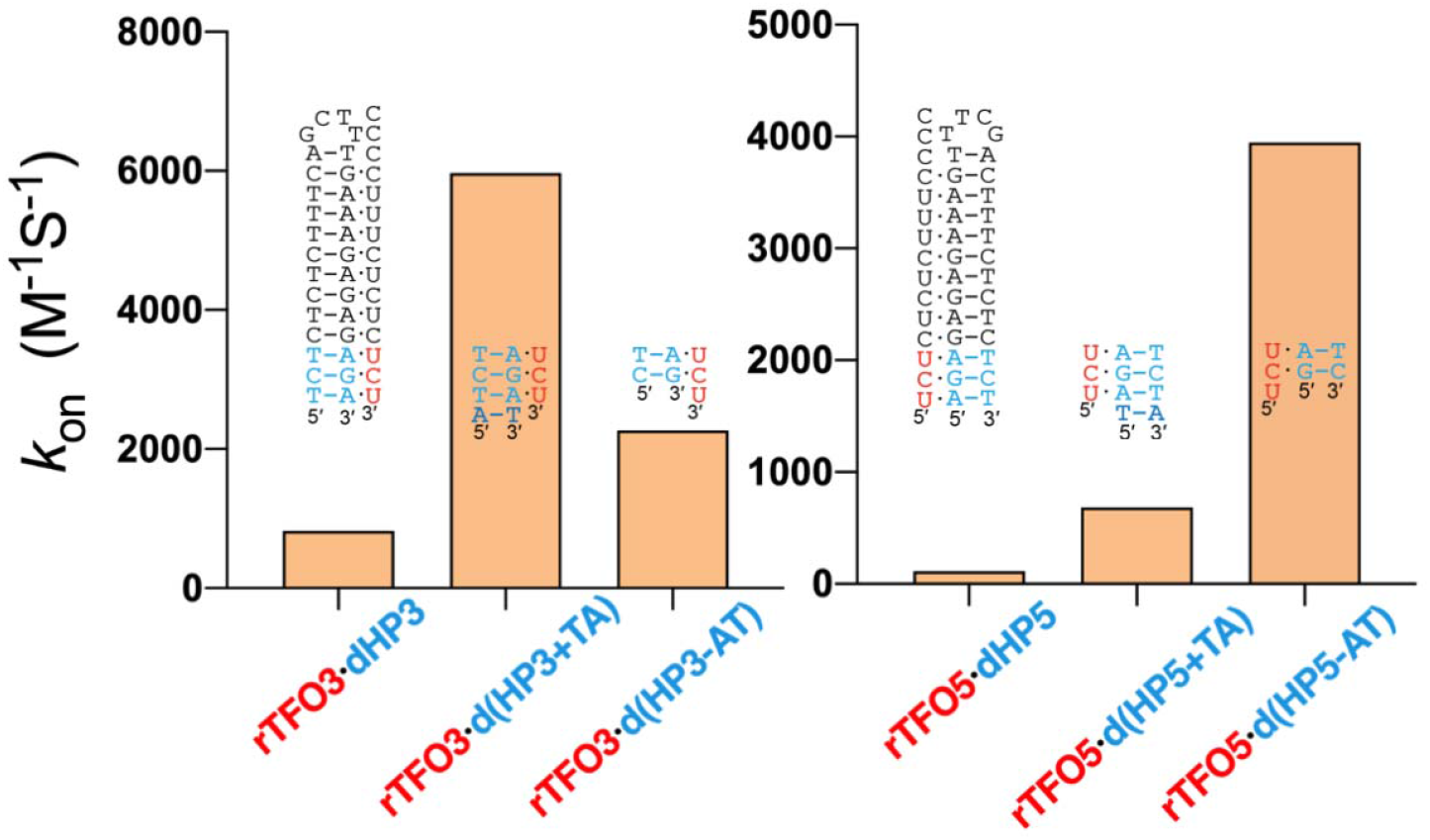

**Highlights:**

- Dangling ends at both third strand and duplex termini enhance triplex stability by accelerating association
- Triplex formation can be nucleated from either end with a bimodal association mechanism
- Terminal blunt-end base triples are dynamic and contribute minimally to stability compared to tailored overhangs

## INTRODUCTION

Nucleic acid triplexes formed within DNA, within RNA, and between DNA and RNA are emerging as important structural elements involved in gene regulation, catalytic functions, and nucleic acid technologies (1-11). A parallel major-groove triplex structure contains multiple pyrimidine·purine-pyrimidine triples (with the “·” and “-” representing Hoogsteen and Watson-Crick pairs, respectively, see **Figure 1**), with the Hoogsteen strand parallel to the purine-rich strand (also denoted as Crick strand) (12-16). The natural parallel major-groove RNA·RNA-RNA triplex structures with consecutive C^+^·G-C and U·A-U major-groove base triples (**Figure 1A**) formed in telomerase RNA pseudoknot are highly conserved and important for telomeric DNA synthesis (17-19). RNA·RNA-RNA triplex formation is also involved in many other functions including intron splicing, translational reading frame recoding, ribozyme catalysis, and riboswitch functions (8-10,20-25). RNA·DNA-DNA triplexes (**Figure 1B**) may form between DNA duplexes and non-coding RNAs including long non-coding RNAs (lncRNAs) and microRNAs facilitating RNA TFO-guided functional regulations of DNA and histones (7,10,11,26-29).. Intramolecular DNA·DNA-DNA triplexes (**Figure 1C**) may also form and are implicated in various biological processes such as chromatin accessibility, DNA replication and transcription, and genomic stability maintenance (6,7,10).

**Figure 1.**
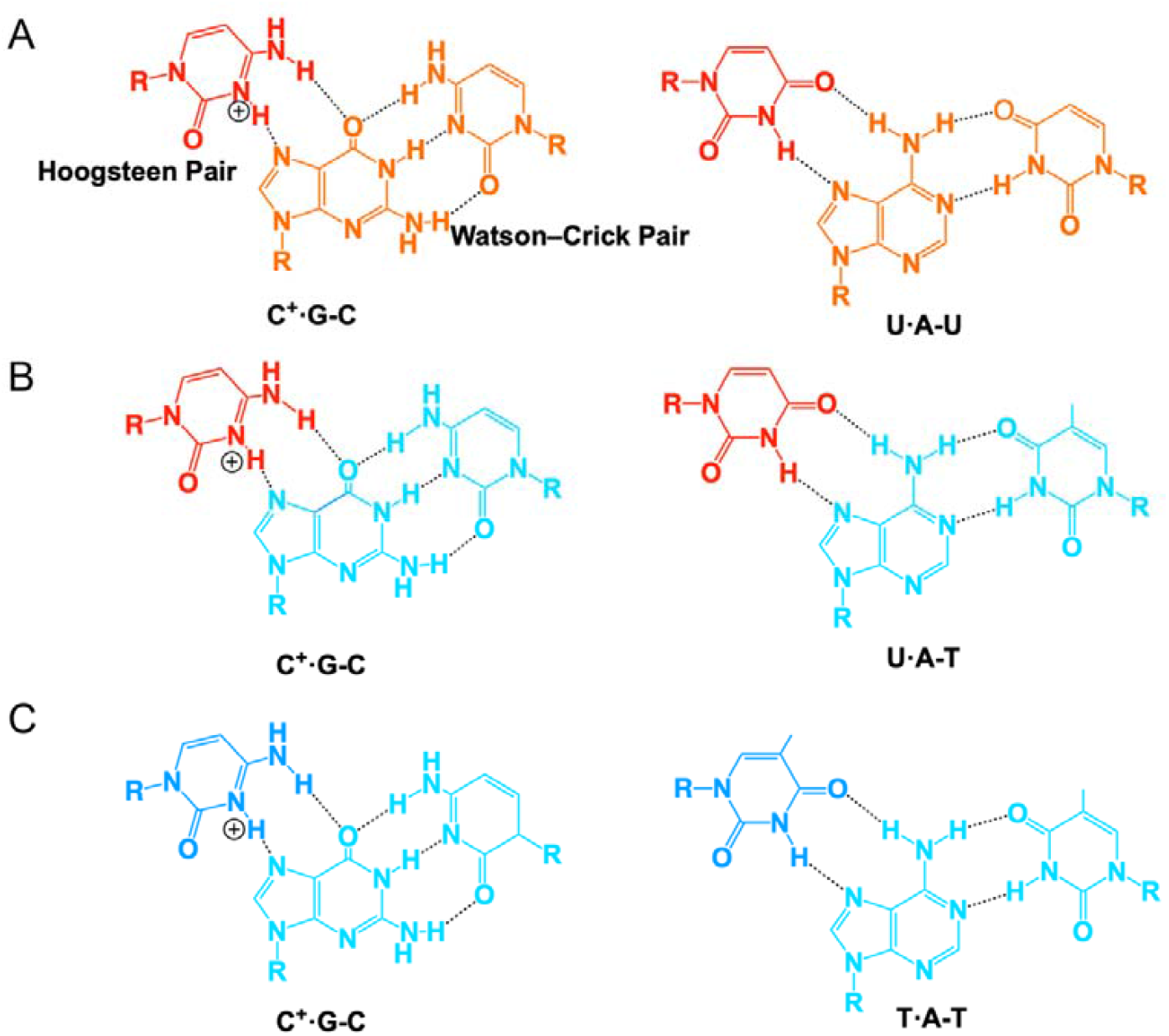
Base triple structures. (A) RNA·RNA-RNA base triples C^+^**·**G-C and U**·**A-U. (B) RNA·DNA-DNA base triples C^+^**·**G-C and U**·**A-T. (C) DNA·DNA-DNA base triples C^+^**·**G-C and T**·**A-T.

Triplex-forming oligonucleotides (TFOs) can be natural nucleic acids as discussed above or artificial nucleic acids that typically bind to the major groove of double-stranded DNA (dsDNA) or dsRNA. A quantitative understanding of the factors affecting triplex formation is needed to facilitate the accurate prediction of natural triplexes and developing triplex-forming ligands targeting dsDNAs and dsRNAs. The effects of nucleic acid composition, sequence, and length on the triplex formation stability have been extensively studied (30-44). Structural context may affect a parallel major-groove triplex formation. For example, a major-groove triplex is often found to be present within an RNA pseudoknot structure (18-20,22,23).

Despite the established thermodynamic stability hierarchy (RNA•DNA-DNA > DNA•DNA-DNA > RNA•RNA-RNA) (30,42,45-48), the kinetic mechanisms governing triplex formation remain largely unexplored. A critical knowledge gap is the influence of terminal non-base-triple-forming nucleotides, or dangling ends, on triplex kinetics. It is thus important to understand how the neighboring sequences and structures (see **Figure 2** for example) affect the nucleic acid triplex stabilities (14,45-53).

**Figure 2:**
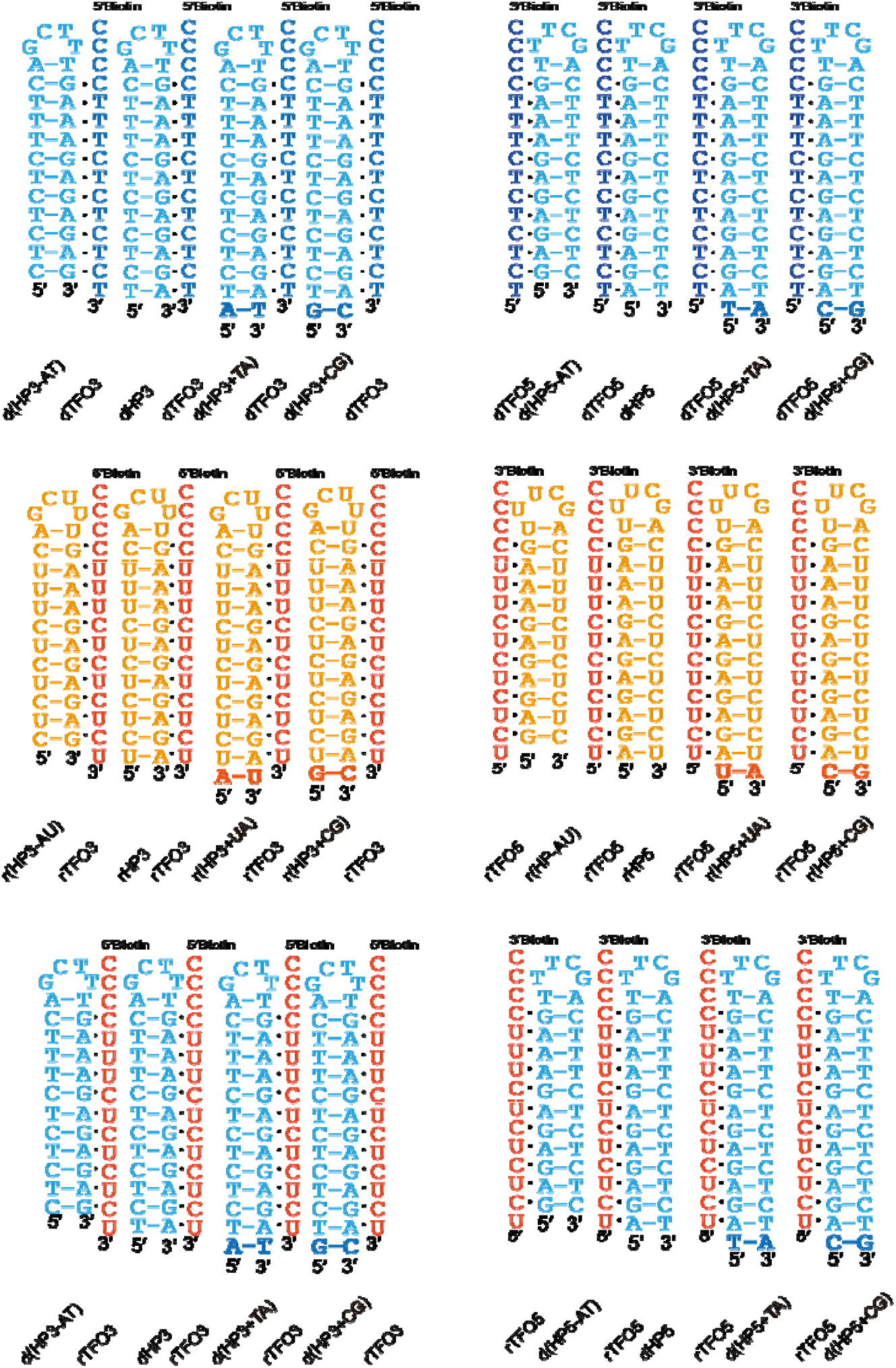
Design of the sequences. Dark blue represents DNA TFOs (dTFO), dark orange represents RNA TFOs (rTFO), light blue represents DNA double strands (dHP), and light orange represents RNA double strands (rHP). All of the triplexes with three different compositions (dTFO•dHP, rTFO•rHP and rTFO•dHP are shown.

The stabilizing effect of dangling nucleotides on duplex stability through stacking interactions is well-established (54-56). However, whether this principle extends to triple helices, and through what kinetic mechanisms, is unknown. Understanding these terminal effects is crucial, as natural triplexes often exist within larger structural contexts with non-canonical termini (23,57).

Here, we utilize bio-layer interferometry (BLI) and circular dichroism (CD) spectroscopy to investigate how terminal residues affect triplex kinetics and stability. We confirm the established stability hierarchy with the BLI-based kinetic methods and demonstrate that both third-strand and duplex dangling ends dramatically enhance triplex stability by accelerating association rates. Our findings support a bimodal association mechanism where triplex formation can be nucleated from either terminus, providing new design principles for triplex-based technologies.

## MATERIALS AND METHODS

### Reagents and materials

All the oligonucleotides used in this study were synthesized and HPLC-purified by Sangon Biotechnology Co., Ltd. (Shanghai, China), and the sequences are listed in Table S1.

#### Reagent

NaCl, MES, and EDTA were obtained from Sangon Biotech (Shanghai) Co., Ltd. MgCl_2_ was from Beyotime Biotechnology (China). All chemical reagents were of analytical grade and used without further purification.

#### Buffer

200 mM NaCl, 20 mM MES, 10 mM MgCl_2_, 0.5 mM EDTA, pH 6.5.

#### Instrument Information

Bio-layer interferometry (BLI) measurement of kinetic parameters was done using Gator® Prime from Gator Bio (Shanghai, China). CD spectrometer was from Applied Photophysics Ltd (England).

#### Bio-layer interferometry assay(BLI)

Using the Streptavidin SA Probe (Gator), the binding kinetics and affinity of immobilized TFOs to various hairpin sequences were determined. Experiments and data analysis were performed using the GatorPrime System (Gator). TFO samples, biotin-labelled at either 3’ or 5’, were diluted by binding buffer (comprising 200 mM NaCl, 20 mM MES, 10 mM MgCl_2_, 0.5 mM EDTA, pH 6.5) to a final concentration of 200nM. Hairpins were diluted with the same solution to different concentrations. The binding and dissociation times were both recorded as 300 seconds at 30°C.

The data were then fitted using Gator Prime software, and the least squares regression was used to model the association followed by dissociation, with the confidence interval (CI) set at 95%. Data were fitted to a 1:1 binding model using Gator Prime software.

### Circular Dichroism assay (CD)

CD experiments were performed in a CD spectrometer using an incubation buffer of 200 mM NaCl, 20 mM MES, 10 mM MgCl_2_, and 0.5 mM EDTA, pH 6.5. 1 μM HP and 1 μM TFO samples were heated and annealed at 95 □ for 5 min, and then rapidly cooled at 4 □ for 10 min. CD spectra were acquired at 30 □, 35 □, and 40 □.

## RESULTS AND DISCUSSION

### Design of the sequences

The design of the study is outlined in **Figure 2**. In our design, we investigated TFOs (rTFO5/dTFO5 versus rTFO3/dTFO3) binding to hairpins from both the 5’ and 3’ ends (rHP5/dHP5 versus rHP3/dHP3). The hairpin sequences rHP5/dHP5 and rHP3/dHP3 form 12-base-triple blunt-ended triplexes with the 12-nucleotide TFOs (rTFO5/dTFO5 and rTFO3/dTFO3, respectively) (see **Figure 2**). On the basis of these hairpins, d(HP5+TA), r(HP5+UA, d(HP5+CG), r(HP5+CG) were designed to add a 5’ dangling base pair in the duplex. Similarly, d(HP3+TA), r(HP3+UA, d(HP3+CG), r(HP3+CG) were designed to add a 3’ dangling base pair in the duplex. With the designed hairpins in hand, we aimed to ascertain whether the presence of an additional dangling base pair impacts the efficiency of triplex formation. Furthermore, we removed the terminal A-U/A-T pair in the hairpins dHP5/rHP5 to generate d(HP5-AT) and r(HP5-AU), to study the contributions of (i) a 5’ blunt-ended base triple and (ii) a 5’ dangling end in TFO. Finally, we removed the terminal A-U/A-T pair in the hairpins dHP3/rHP3 to generate d(HP3-AT) and r(HP3-AU), to study the contributions of (i) a 3’ blunt-ended base triple and (ii) a 3’ dangling end in TFO.

### BLI Validation of Triplex Stability Hierarchy: RDD > DDD > RRR > DRR

Our BLI analysis quantitatively confirmed the established triplex stability hierarchy. The equilibrium dissociation constants (*K*_D_) based on the BLI assay revealed RNA•DNA-DNA (RDD, rTFO•dHP) as the most stable (*K*_D_ = 7.48 μM for rTFO5•dHP5), followed by DNA•DNA-DNA (DDD, dTFO•dHP, *K*_D_ = 14.2 μM for for dTFO5•dHP5), and RNA•RNA-RNA (RRR, rTFO•rHP, *K*_D_ = 38.1 μM for rTFO5•rHP5). DNA•RNA-RNA (DRR) showed no measurable binding, consistent with previous reports (**Table 1 and Figures 3, 4, Supplementary Figure S1**) (30-32,42). The NMR and crystal structures of various triplexes reveal that the base-base hydrogen bonds (**Figure 1**) are essentially the same for the three types of triplexes (18,19,22,23,47,48,58,59). The different helical shapes of DNA and RNA duplexes are expected to result in variations of the stacking of the base triples and backbone-backbone interactions. It is likely that, compared to an RNA duplex, a DNA duplex with a less twisted helical structure, wider major groove, and lower density of negatively charged phosphate groups, is more favorable for the formation of a major-groove triplex with an RNA or DNA TFO (Hoogsteen strand).

**Table 1.** BLI results of TFOs binding to hairpins.

| BLI results |  |  |  |  |
| --- | --- | --- | --- | --- |
| *Name* | $K_D$ ($\mu\text{M}$) | $k_{\text{on}}$ ($\text{M}^{-1} \text{s}^{-1}$) | $k_{\text{off}}$ ($10^{-6} \text{s}^{-1}$) | $R^2$ |
| dTFO5•dHP5 | 14.2±0.18 | 218±2 | 3090±4 | 0.99 |
| dTFO5•d(HP5+TA) | 2.33±0.01 | 946±4 | 2210±3 | 0.99 |
| dTFO5•d(HP5+GC) | 4.14±0.04 | 596±6 | 2470±6 | 0.99 |
| dTFO5•d(HP5-AT) | 3.84±0.01 | 1270±7 | 1170±12 | 0.99 |
| rTFO5•dHP5 | 7.48±0.21 | 114±6 | 754±4 | 0.99 |
| rTFO5•d(HP5+TA) | 0.73±0.01 | 686±4 | 501±3 | 0.99 |
| rTFO5•d(HP5+GC) | 1.01±0.01 | 748±4 | 757±5 | 0.99 |
| rTFO5•d(HP5-AT) | 0.47±0.01 | 3950±18 | 1760±5 | 0.99 |
| rTFO5•rHP5 | 38.1±0.92 | 111±20 | 4240±13 | 0.98 |
| rTFO5•r(HP5+UA) | 4.48±0.07 | 1040±14 | 4658±14 | 0.97 |
| rTFO5•r(HP5+GC) | 7.49±0.1 | 294±1 | 2200±1 | 0.98 |
| rTFO5•r(HP5-AU) | 6.54±0.07 | 237±2 | 1550±2 | 0.99 |
| dTFO5•rHP5 |  | No Binding |  |  |
| dTFO5•r(HP5+UA) |  | No Binding |  |  |
| dTFO5•r(HP5+GC) |  | No Binding |  |  |
| dTFO5•r(HP5-AU) |  | No Binding |  |  |
| dTFO3•dHP3 | 474 | 3 | 1290±7 | 0.99 |
| dTFO3•d(HP3+TA) | 3.26±0.03 | 1880±14 | 6140±13 | 0.98 |
| dTFO3•d(HP3+CG) | 20±1.90 | 1120±10 | 22500±52 | 0.99 |
| dTFO3•d(HP3-AT) | 1.84±0.02 | 774±4 | 1430±3 | 0.99 |
| rTFO3•dHP3 | 8.96±0.17 | 821±15 | 7360±19 | 0.96 |
| rTFO3•d(HP3+TA) | 0.70±0.01 | 5970±31 | 4160±9 | 0.99 |
| rTFO3•d(HP3+CG) | 1.42±0.01 | 3080±14 | 4360±7 | 0.98 |
| rTFO3•d(HP3-AT) | 0.78±0.02 | 2269±50 | 1770±11 | 0.99 |
| rTFO3•rHP3 | 3930 | 5 | 20500±128 | 0.95 |
| rTFO3•r(HP3+UA) | 4.57±0.04 | 1500±11 | 6840±12 | 0.99 |
| rTFO3•r(HP3+CG) | 15.7±0.87 | 966±53 | 15100±99 | 0.92 |
| rTFO3•r(HP3-AU) | 5.92±0.03 | 663±3 | 3930±4 | 0.98 |
| dTFO3•rHP3 |  | No Binding |  |  |
| dTFO3•r(HP3+UA) |  | No Binding |  |  |
| dTFO3•r(HP3+CG) |  | No Binding |  |  |
| dTFO3•r(HP3-AU) |  | No Binding |  |  |

**Figure 3:**
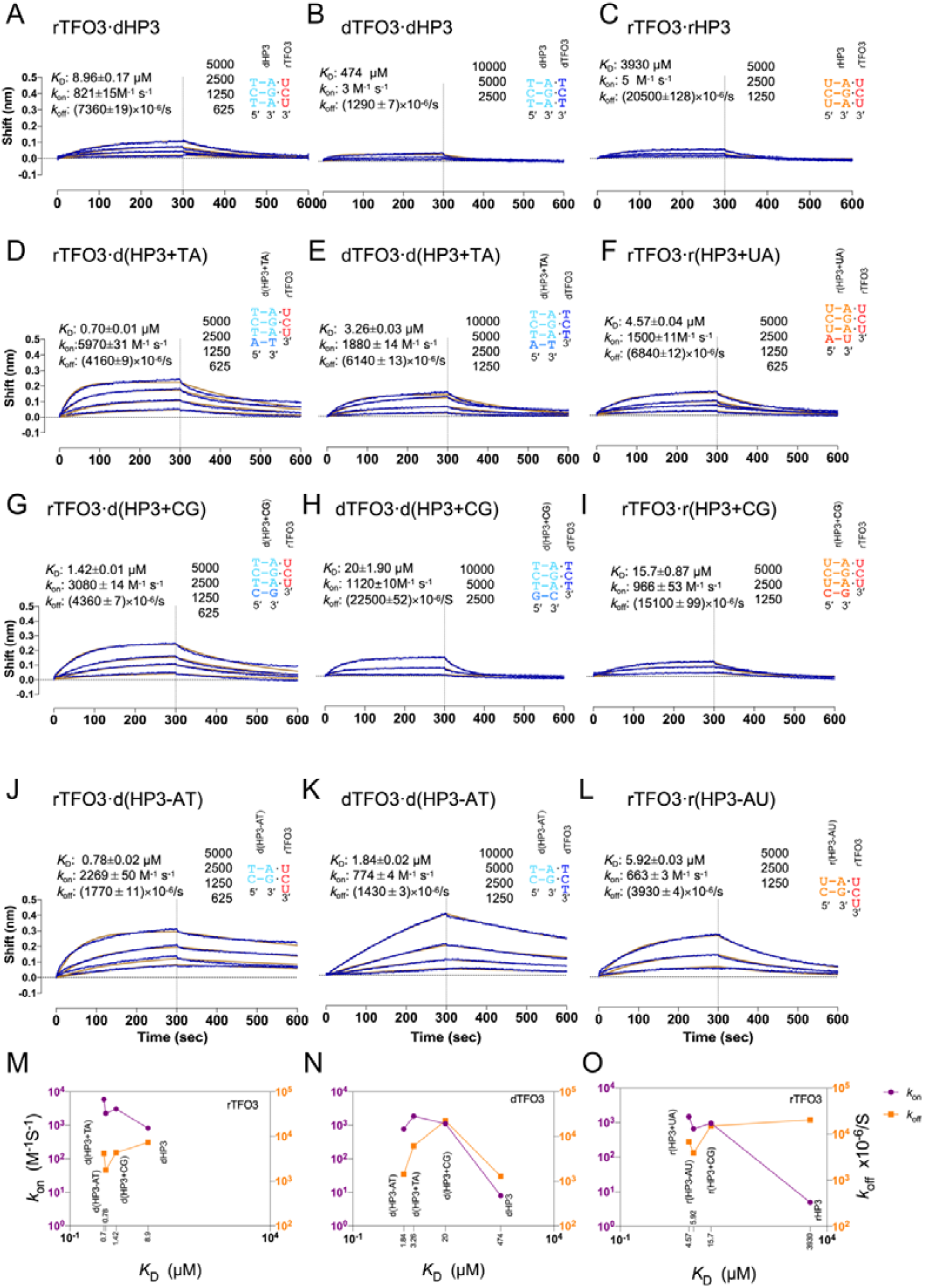
BLI results of TFO3 binding to hairpins with modifications at the 3’ ends for studying the effects of 3’ dangling ends. (A, D, G, J, M) BLI data and plot of binding kinetics versus *K*_D_ for rTFO3 binding to dHP3 and modified hairpins. (B, E, H, K, N) BLI data and plot of binding kinetics versus *K*_D_ for dTFO3 binding to dHP3 and modified hairpins., (C,F,I,L,O) BLI data and plot of binding kinetics versus *K*_D_ for rTFO3 binding to rHP3 and modified hairpins. The assay was done at 30 °C using a buffer of 200 mM NaCl, 20 mM MES, 10 mM MgCl_2_, 0.5 mM EDTA, pH 6.5.

**Figure 4:**
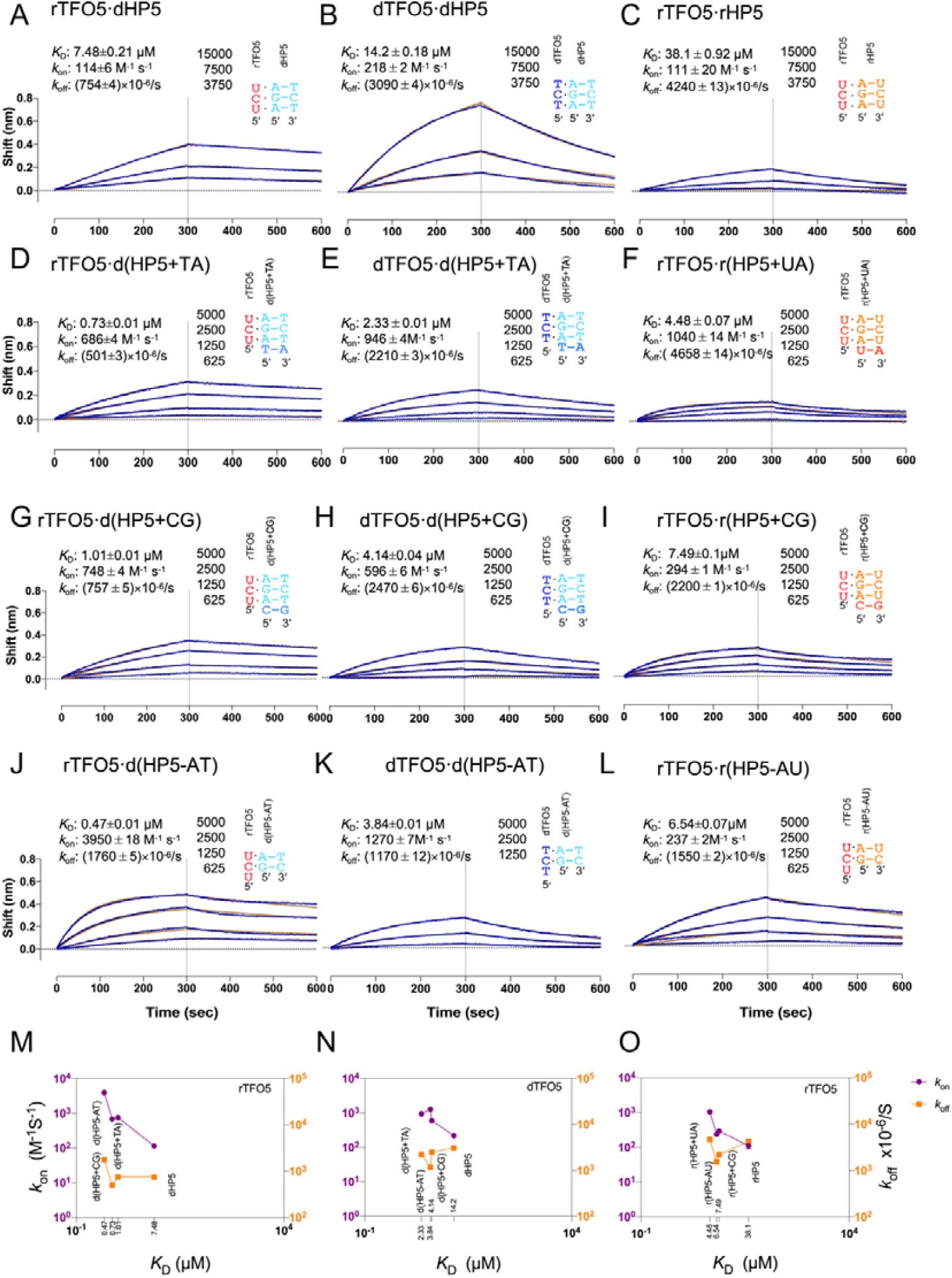
BLI results of TFO5 binding to hairpins with modifications at the 5’ ends for studying the effects of 5’ dangling ends. (A, D, G, J, M) BLI data and plot of binding kinetics versus *K*_D_ for rTFO5 binding to dHP5 and modified hairpins. (B, E, H, K, N) BLI data and plot of binding kinetics versus *K*_D_ for dTFO5 binding to dHP5 and modified hairpins., (C,F,I,L,O) BLI data and plot of binding kinetics versus *K*_D_ for rTFO5 binding to rHP5 and modified hairpins. The assay was done at 30 °C using a buffer of 200 mM NaCl, 20 mM MES, 10 mM MgCl_2_, 0.5 mM EDTA, pH 6.5.

### Both 3’ and 5’ Dangling Ends of the Duplex Drive Stabilization through Enhanced Association

We further tested whether introducing a dangling base pair into the hairpins affects triplex formation. The BLI experimental results show that introducing a dangling base pair at the 3’ terminus effectively promotes the triplex formation between TFOs and hairpins (with smaller *K*_D_ values, **Table 1, Figure 3A-I**). Kinetic dissection revealed this was primarily driven by a substantial increase in the association rate (*k*_on_) (**Table 1, Figure 3M-O**), suggesting the duplex dangling end acts as a pre-organized platform that facilitates the initial docking of the TFO.

Crucially, a similar stabilizing effect was observed upon extending the duplex with a dangling base pair at the 5’ end of rTFO5/dTFO5. For instance, the binding of rTFO5 to hairpins r(HP5+UA/CG) and d(HP5+TA/CG) showed an enhanced affinity and a faster *k*_on_ compared to the blunt-ended controls rHP5 and dHP5 (**Table 1, Figure 4A-I, M-O**).

The fact that extending either the 5’ or the 3’ duplex dangling end enhances *k*_on_ challenges a model where triplex formation initiates exclusively from one terminus (60) (61). Instead, it supports a mechanism where the binding can be initiated from both ends, with dangling duplex base pair at either location providing a favorable stacking interface to nucleate triple-helix formation. The controversy may be due to the possibility that replacing a terminal base triple with a mismatached triple may not weaken triplex formation (60). Thus, strategic modifications to both of the duplex termini dramatically enhanced triplex stability. This enhancement through duplex dangling ends (overhangs) demonstrates that the duplex dangling end serves as a critical nucleation platform that facilitates initial TFO docking. The consistent acceleration of *k*□□ across different triplex types indicates this is a general mechanism for triplex stabilization.

Concurrently, circular dichroism (CD) spectra at different temperatures reveal that the triplex structure with blunt ends weakens significantly with increasing temperature (**Supplementary Figures S2, S3**). The characteristic peak near 210 nm gradually diminishes as the temperature rises, with essentially no peak at 35 °C and 40 °C. However, the CD spectra of rTFO5/rTFO3 with d(HP5+TA)/d(HP3+TA) exhibit a characteristic peak near 210 nm even at 35 °C and 40 °C. These findings indicate that the triplex structure stability is enhanced by adding a dangling base pair in the duplex.

### Both 3’ and 5’ Dangling Ends in the TFO Drive Stabilization through Enhanced Association

We next investigated whether extending the single-stranded TFO to generate a dangling base affects triplex formation. The sequences were designed by removing a terminal base pair in the hairpins to generate d(HP3-AT), r(HP3-AU), d(HP5-AT), and r(HP5-AU). The hairpins are expected to form 11 base triples (instead of 12 base triples with a blunt end) with a dangling TFO base. BLI assay data (see **Figures 3 and 4, Table 1**) suggest that removal of a base pair at both ends results in a significantly enhanced triplex formation, as evidenced by markedly reduced *K*_D_ values, increased *k*_on_ values and decreased *k*_off_ values (**Table 1, Figure 3A–C, J–L, M–O and Figure 4A–C, J–L, M–O**). For example, creating a single-base dangling end (overhang) at the duplex 5’ end consistently improved binding affinity and kinetics. For RNA•DNA-DNA triplexes, rTFO5 binding to d(HP5-AT) (*K*_D_ = 0.47 μM, *k*□□ = 3950 M□^1^s□^1^, **Figure 4J**) was significantly stronger and faster than to blunt-ended dHP5 (*K*_D_ = 7.48 μM, *k*□□ = 114 M□^1^s□^1^, **Figure 4A**), representing a 16-fold increase in affinity and a 35-fold acceleration in association. Similarly, for DNA•DNA-DNA triplexes, dTFO5 binding to d(HP5-AT) (*K*_D_ = 3.84 μM, *k*□□ = 1270 M□^1^s□^1^, **Figure 4K**) substantially outperformed binding to dHP5 (*K*_D_ = 14.2 μM, *k*□□ = 218 M□^1^s□^1^, **Figure 4B**).

Similarly, rTFO3 binding to d(HP3-AT) (*K*_D_ = 0.78 μM, *k*□□ = 2269 M□^1^sL^1^, **Figure 3J**) was significantly stronger and faster than to blunt-ended dHP3 (*K*_D_ = 8.96 μM, *k*□□ = 821 ML^1^sL^1^, **Figure 3A**). Furthermore, dTFO3 binding to d(HP3-AT) (*K*_D_ = 1.84 μM, *k*□□ = 774 ML^1^sL^1^, **Figure 3K**) shows significantly stronger binding to dHP3 (*K*_D_ = 474 μM, *k*□□ = 3 ML^1^□L^1^, **Figure 3B**).

### Bimodal Association from Both TFO Strand and Duplex Ends

The complementary stabilization observed from both duplex dangling ends points to a bimodal association mechanism. Triplex formation can be nucleated from either end of the binding interface, with dangling residues at either location providing favorable stacking interfaces.

The base triple stacking patterns from the reported triplex structures (**Figure 5**) reveal that the 5′ terminal residues of the TFOs have varied stacking with the next base triple, with an RNA·RNA-RNA triplex having more intra-strand stacking interactions within the TFO, and DNA duplex-containing triplexes potentially having cross-strand stacking between the TFO and Crick strand triplexes (18,19,22,23,47,48,58,59). The results presented herein suggest that the potentially more distorted B-type DNA double strand may permit the dangling Watson-Crick DNA base pairs to interact with the 5′-end bases in the TFO for cross-strand stacking (31,38,47,58). It is expected that the dangling base pair in the duplex and dangling base in the TFO may exhibit local distortion upon triplex formation **Figure 5** (47). We may also expect that terminal mismatches may contribute significantly to triplex formation through stacking and hydrogen bonding interactions (**Figure 5**). The results obtained from this study indicate that the major-groove parallel triplex with DNA or RNA Watson-Crick duplexes can be substantially stabilized by strategic extension of either the Watson-Crick duplex or the TFO at both the 5′ and 3′ ends.

**Figure 5.**
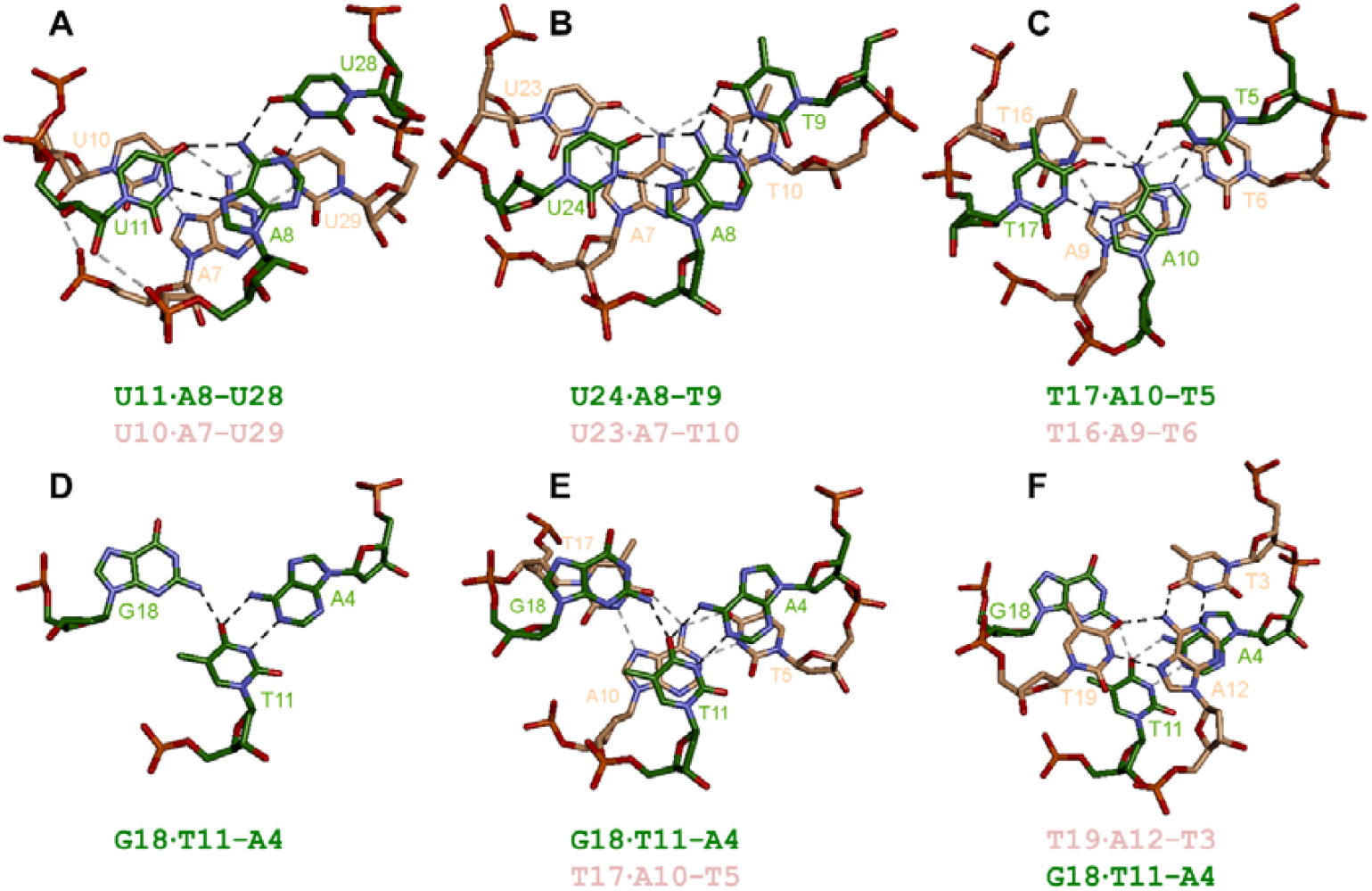
Hydrogen bonding and stacking of base triples. (A) Two RNA·RNA-RNA U·A-U base-triples (PDB ID: 3P22 (22)). There is a backbone-backbone hydrogen bond formed between 2’-OH in TFO and phosphate oxygen in purine strand (indicated with gray dashed lines). (B) Two RNA·DNA-DNA U·A-T base-triples (PDB ID: 1R3X (58)). The backbone-backbone hydrogen bond between the 2’-OH in the TFO and a phosphate oxygen in the purine strand is not present in the NMR structure (58), although it was previously predicted to form (31)). (C) Two DNA·DNA-DNA T·A-T base-triples (PDB ID: 149D (59)). (D-F) A DNA·DNA-DNA G·T-A base-triple and the stacking involving the G·T-A triple (PDB ID: 149D (59)). The G residue in the TFO has stacking with the 5’ side residue (panel E) but no stacking with the 3’ side residue (panel F).

## CONCLUSIONS

In this study, our BLI analysis quantitatively confirmed the established triplex stability hierarchy (RNA•DNA-DNA > DNA•DNA-DNA > RNA•RNA-RNA, with no DNA•RNA-RNA triplex forming). Our kinetic study reveals that triplex stability is governed by dynamic terminal interactions rather than simply the number of base triples. We demonstrate that dangling ends from both the TFO strand and duplex dramatically enhance triplex stability by accelerating association rates, supporting a bimodal nucleation mechanism. Specifically, both the addition of a base pair to the duplex (e.g., creating d(HP5+TA)) and the removal of a terminal base pair from the duplex (e.g., creating d(HP5-AT)) to create tailored dangling ends (overhangs) produce more stable triplexes than their blunt-ended counterparts. This symmetrically underscores the minimal contribution of dynamic terminal triples and the critical role of terminal stacking. These insights establish that optimal triplex design requires strategic optimization of both TFO and duplex terminal structures, providing a new, general design principle for developing effective nucleic acid triplex-based technologies in gene regulation, chromatin engineering, and nanotechnology (62-65).

## Author contributions

Conceptualization, Methodology, Writing - original draft preparation: Fangyu Zhou, Yiran Liu, Gang Chen; Formal analysis and investigation: Gang Chen, Fangyu Zhou; Resources: Fangyu Zhou, Gang Chen, Yongqiang Wang; Writing – Review & Editing.: Fangyu Zhou, Gang Chen; Funding acquisition, Supervision: Gang Chen, Yongqiang Wang.

## Data availability

The authors confirm that the data presented in this article are available upon request.

