## Supplementary material for "Dangling Ends of Third Strand and Duplex Drive Nucleic Acid Triplex Stabilization through Bimodal Association": SI

Table S1. The sequences used in this study.

| Sequences name | Nucleic acid type | Sequence (5'-3') |
| --- | --- | --- |
| dTFO5 | DNA | TCTCTCTCTTTCCCC-Biotin |
| rTFO5 | RNA | UCUCUCUCUUUCCCC-Biotin |
| dHP5 | DNA | AGAGAGAGAAAGTTTCGACTTTCTCTCTCT |
| d(HP5-AT) | DNA | GAGAGAGAAAGTTTCGACTTTCTCTCTC |
| d(HP5+TA) | DNA | TAGAGAGAGAAAGTTTCGACTTTCTCTCTCTA |
| d(HP5+CG) | DNA | CAGAGAGAGAAAGTTTCGACTTTCTCTCTCTG |
| rHP5 | RNA | AGAGAGAGAAAGUUUCGACUUUCUCUCUCU |
| r(HP5-AU) | RNA | GAGAGAGAAAGUUUCGACUUUCUCUCUC |
| r(HP5+UA) | RNA | UAGAGAGAGAAAGUUUCGACUUUCUCUCUCUA |
| r(HP5+CG) | RNA | CAGAGAGAGAAAGUUUCGACUUUCUCUCUCUG |
| dTFO3 | DNA | Biotin-CCCCTTTCTCTCTCT |
| rTFO3 | RNA | Biotin-CCCCUUUCUCUCUCU |
| dHP3 | DNA | TCTCTCTCTTTCAGCTTTGAAAGAGAGAGA |
| d(HP3-AT) | DNA | CTCTCTCTTTCAGCTTTGAAAGAGAGAG |
| d(HP3+TA) | DNA | ATCTCTCTCTTTCAGCTTTGAAAGAGAGAGAT |
| d(HP3+CG) | DNA | GTCTCTCTCTTTCAGCTTTGAAAGAGAGAGAC |
| rHP3 | RNA | UCUCUCUCUUUCAGCUUUGAAAGAGAGAGA |
| r(HP3-AU) | RNA | CUCUCUCUUUCAGCUUUGAAAGAGAGAG |
| r(HP3+UA) | RNA | AUCUCUCUCUUUCAGCUUUGAAAGAGAGAGAU |
| r(HP3+CG) | RNA | GUCUCUCUCUUUCAGCUUUGAAAGAGAGAGAC |

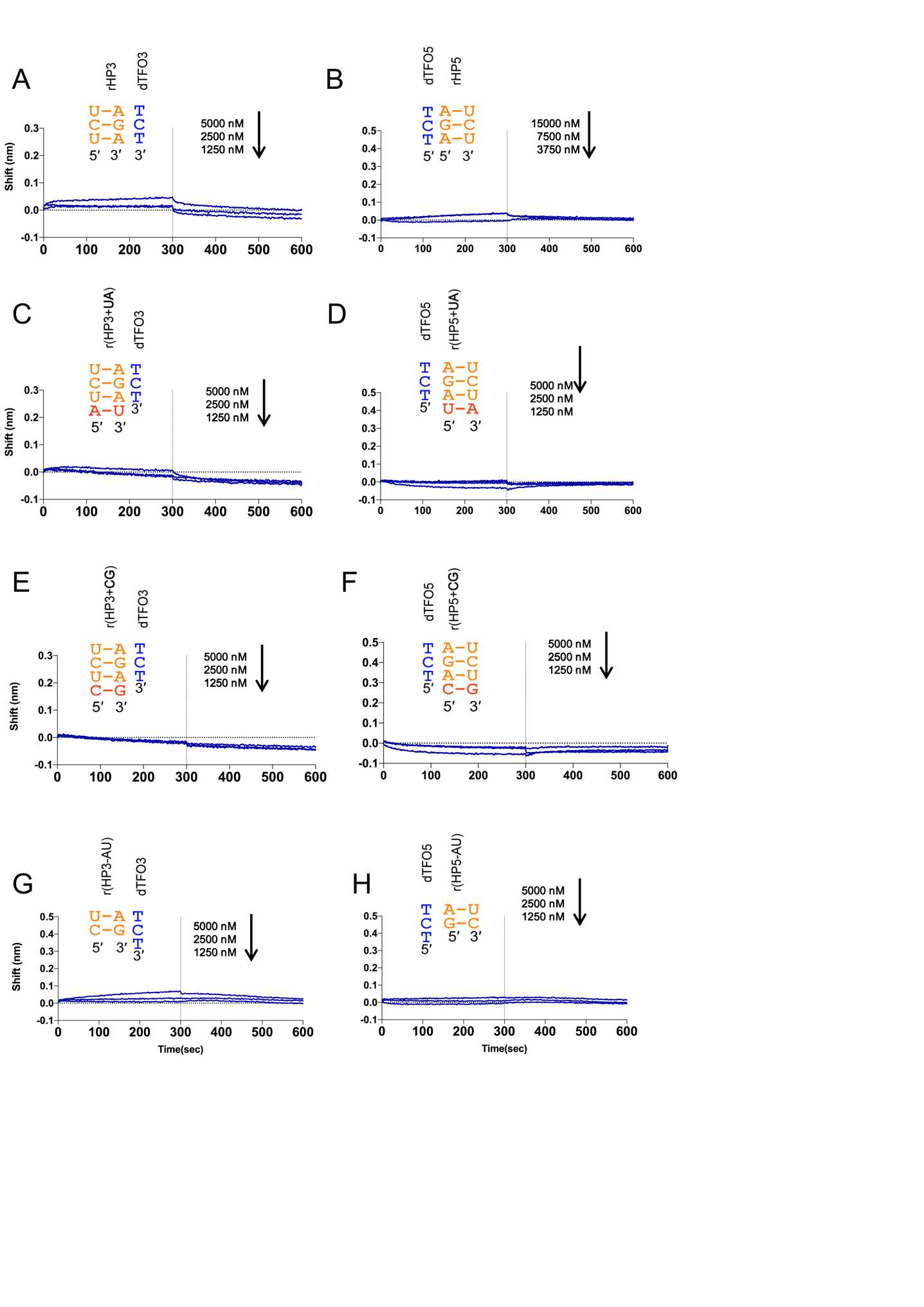

**Figure S1**: No observable binding based on BLI results of dTFO binding to rHP. (A,C,E,G): BLI testing of the binding of dTFO3 to rHP3 and modified hairpins. (B,D,F,H): BLI testing of the binding of dTFO5 binding to rHP5 and modified hairpins. BLI data showing no measurable binding is consistent with the DNA•RNA-RNA (DRR) triplex instability reported in the main text.

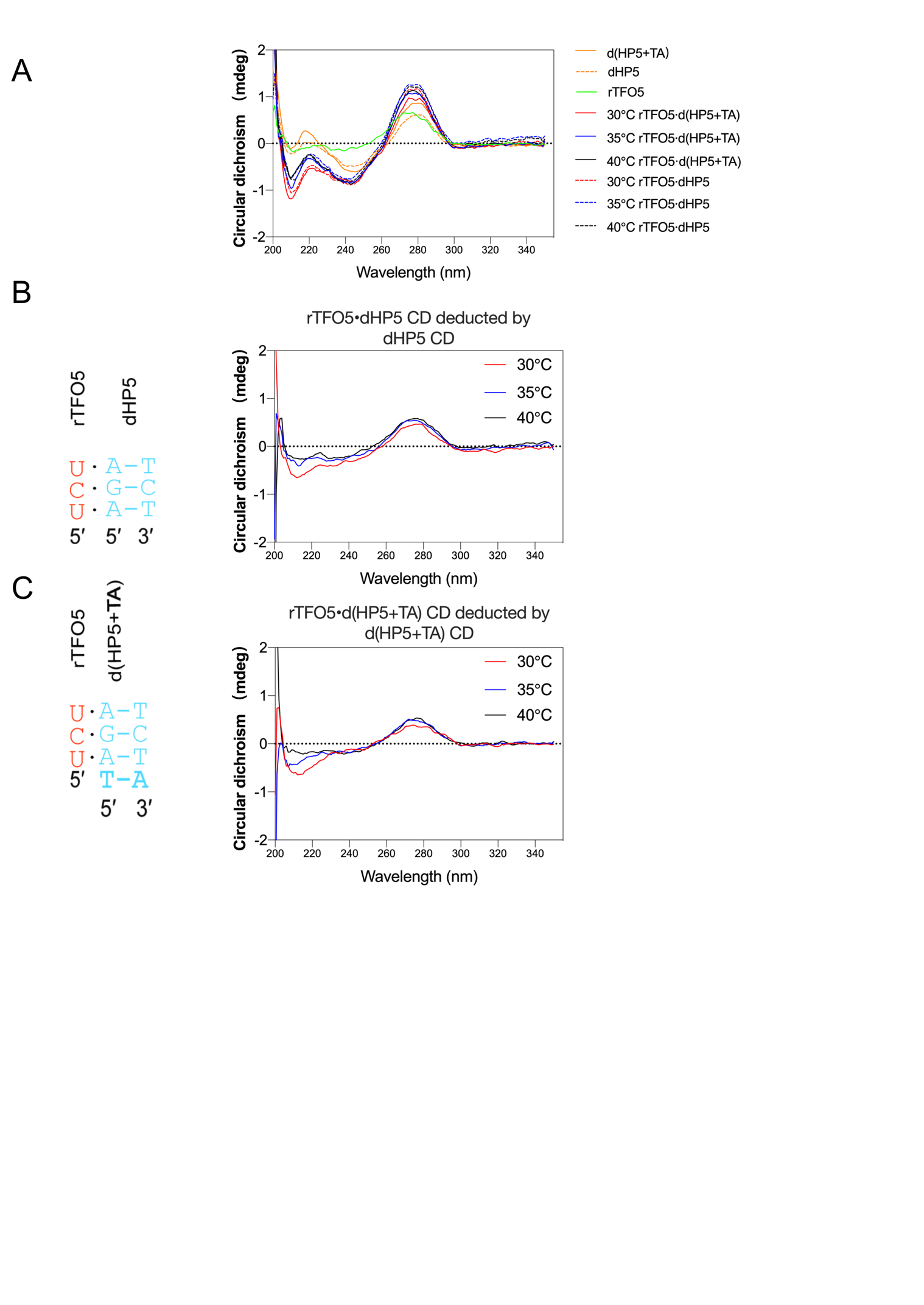

Figure S2: CD spectra of rTFO5 binding to dHP5 and d(HP5+TA). A: CD spectra at varied temperatures. B: Differential CD spectra of rTFO5•dHP5 triplex and dHP5 at different temperatures. C: Differential CD spectra of rTFO5•d(HP5+TA) and d(HP5+TA) at different temperatures.

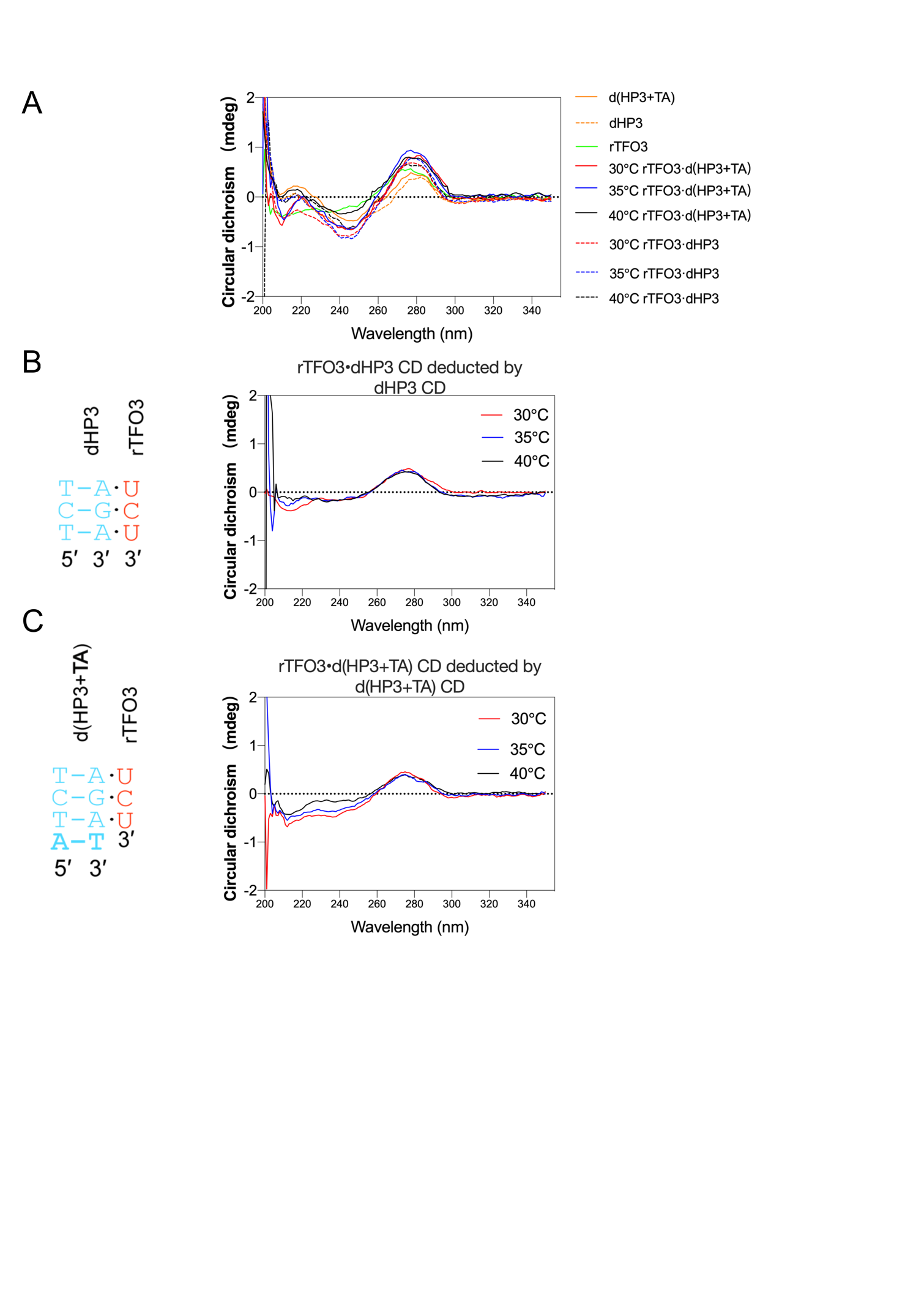

Figure S3: CD spectra of rTFO3 binding to dHP3 and d(HP3+TA). A: CD spectra at varied temperatures. B: Differential CD spectra of rTFO3•dHP3 triplex and dHP3 at different temperatures. C: CD spectra of rTFO3•d(HP3+TA) triplex with d(HP3+TA) at different temperatures.
